# Transcriptional profiling of hepatocytes infected with the replicative form of the malaria parasite *Plasmodium cynomolgi*

**DOI:** 10.1101/2022.07.18.500418

**Authors:** Gabriel Mitchell, Guglielmo Roma, Annemarie Voorberg-van der Wel, Martin Beibel, Anne-Marie Zeeman, Sven Schuierer, Laura Torres, Erika L Flannery, Clemens HM Kocken, Sebastian A. Mikolajczak, Thierry T. Diagana

**Affiliations:** Open Innovation at Novartis Institute for Tropical Diseases, Novartis Institutes for BioMedical Research, Emeryville, California, USA; Chemical Biology & Therapeutics, Novartis Institutes for BioMedical Research, Basel, Switzerland; Department of Parasitology, Biomedical Primate Research Centre, Rijswijk, The Netherlands; Novartis Institute for Tropical Diseases, Novartis Institutes for BioMedical Research, Emeryville, California, USA

**Keywords:** Relapsing malaria, monkey, schizonts, liver stages, host response

## Abstract

The zoonotic simian parasite *Plasmodium cynomolgi* develops into replicating schizonts and dormant hypnozoites during the infection of hepatocytes and is used as a model organism to study relapsing malaria. We previously reported the transcriptional profiling of *P. cynomolgi* liver stages and revealed many important biological features of the parasite (Bertschi et al., Elife, 2018; Voorberg-van der Wel et al., Elife, 2017) but left out the host response to malaria infection. Here, we used our published RNA sequencing data to quantify the expression of host genes in rhesus macaque hepatocytes infected with *P. cynomolgi* in comparison to either cells from uninfected samples or uninfected bystander cells. Although the dataset could not be used to resolve the transcriptional profile of hypnozoite-infected hepatocytes, it provided a snapshot of the host response to liver stage schizonts and identified specific host pathways that are modulated during malaria infection. This study constitutes a valuable resource characterizing the hepatocyte response to *P. cynomolgi* infection and provides a framework to build on future research that aims at understanding hepatocyte-parasite interactions during relapsing malaria infection.

## INTRODUCTION

The protozoan parasites *Plasmodium falciparum* and *Plasmodium vivax* are main sources of human malaria, a disease that globally impacts hundreds of millions of lives each year (WHO, 2021). The lifecycle of malaria includes an asymptomatic liver stage of infection during which the genome from a single parasite is amplified thousands of times in a process known as schizogony, heavily relying on interactions with the host cell (Vaughan & Kappe, 2017). In addition to form replicative liver stage schizonts, *P. vivax* and other relapsing malaria species can differentiate into dormant hypnozoites that can reactivate months or even years after the initial infection, and for which only limited treatment options are available (Schafer et al., 2021). A better understanding of the host processes required during the liver stages of malaria might lead to the identification of targets for the development of host-directed therapies (Wei et al., 2021). Host-directed therapies are especially relevant to hypnozoites considering the tolerance of non-replicating microbes to conventional antimicrobials (Brauner et al., 2016).

Efforts to understand the biology of hypnozoites and to develop radical cure treatments against malaria are dampened by major technical challenges, some of which relate to inherent characteristics of *P. vivax*. In this regard, the zoonotic simian parasite *P. cynomolgi* (Bykersma, 2021) has been used as a model organism to study relapsing malaria and the biology of liver stage hypnozoites (Schafer et al., 2021; Voorberg-van der Wel et al., 2020). Using FACS-purification of hepatocytes infected with GFP-expressing transgenic parasites, our group previously reported the transcriptional profiles of *P. cynomolgi* liver stages and revealed many important biological features of the parasite but did not investigate the host response to malaria infection (Bertschi et al., 2018; Gupta et al., 2019; Voorberg-van der Wel et al., 2017). In this follow-up study, we re-analyzed our previously published RNA sequencing data to define the transcriptional signature of cultured rhesus macaque hepatocytes infected with *P. cynomolgi*.

## RESULTS

### Transcriptomic analysis of primary hepatocytes infected with *P. cynomolgi* schizonts

To gain insight into the host response during the liver stages of malaria, previously published RNA sequencing datasets obtained from primary simian hepatocytes infected with *P. cynomolgi* (Bertschi et al., 2018; Voorberg-van der Wel et al., 2017) were aligned to a reference host genome and used to quantify the expression of host genes in comparison to uninfected cells (see Materials and Methods). More precisely, *P. cynomolgi*-infected hepatocytes were cultured for 6-10 days and subsequently FACS-purified yielding GFP-expressing hypnozoites (GFP-low) and schizonts (GFP-high). These were compared to cells from uninfected samples using transcriptional analysis. Preliminary results showed that while samples infected for 6-7 days were associated with only few statistically significant differentially expressed genes, samples infected for 9-10 days were associated with numerous statistically significant changes in comparison to uninfected samples (data not shown). As GFP-low samples were previously shown to be contaminated with uninfected cells as well as with schizont or released merozoite transcripts at 9-10 day post-infection (dpi) (Bertschi et al., 2018), the GFP-low samples were excluded from this study (See Table 1 for a descriptive list of samples used in this study) and the hepatocyte response to hypnozoites was not determined. Despite this, the 9-10 dpi GFP-high RNAseq samples were further analyzed to provide a snapshot of the transcriptional response of primary rhesus macaque hepatocytes to *P. cynomolgi* schizonts.

**Table 1.**
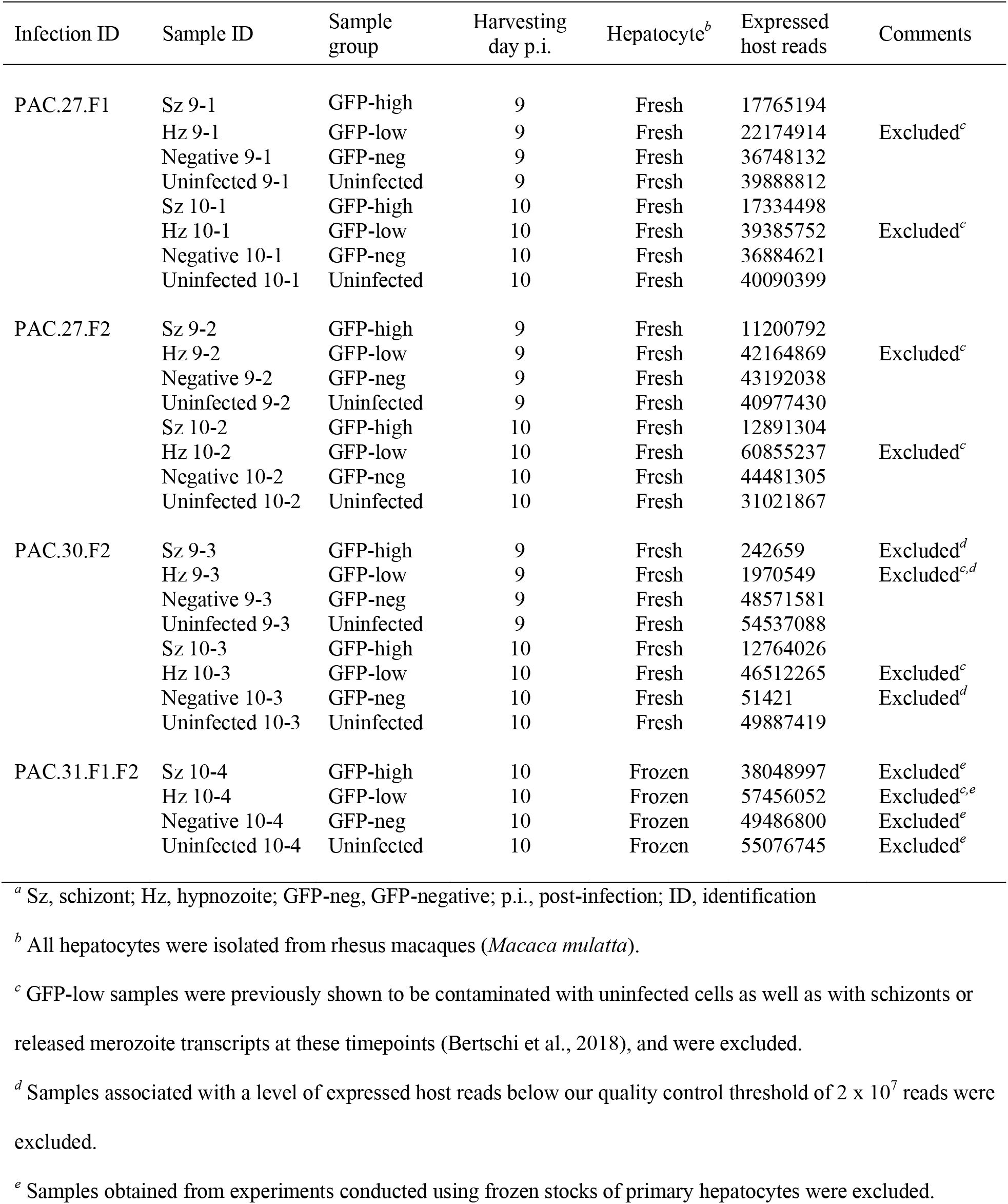
Samples used in this study*^a^*

Global transcriptional differences between schizont-infected cells, uninfected bystander (GFP-negative, negative) cells and cells from uninfected samples were evaluated using sample-to-sample distance analyses (Figures 1A and 1B). While clusters for schizont-infected cells and cells from uninfected samples were clearly separated in space, the clustering of the negative samples were not as defined. It was reasoned that while the comparison of schizont-infected cells to uninfected samples might reveal a larger number of statistically significant gene modulations, the comparison of schizont-infected cells to uninfected bystander cells was likely to be more stringent and reveal genes specifically modulated in schizont-infected hepatocytes. Therefore, the transcriptional response of schizont-infected hepatocytes was determined in comparison to both uninfected samples and uninfected bystander cells.

**Figure 1.**
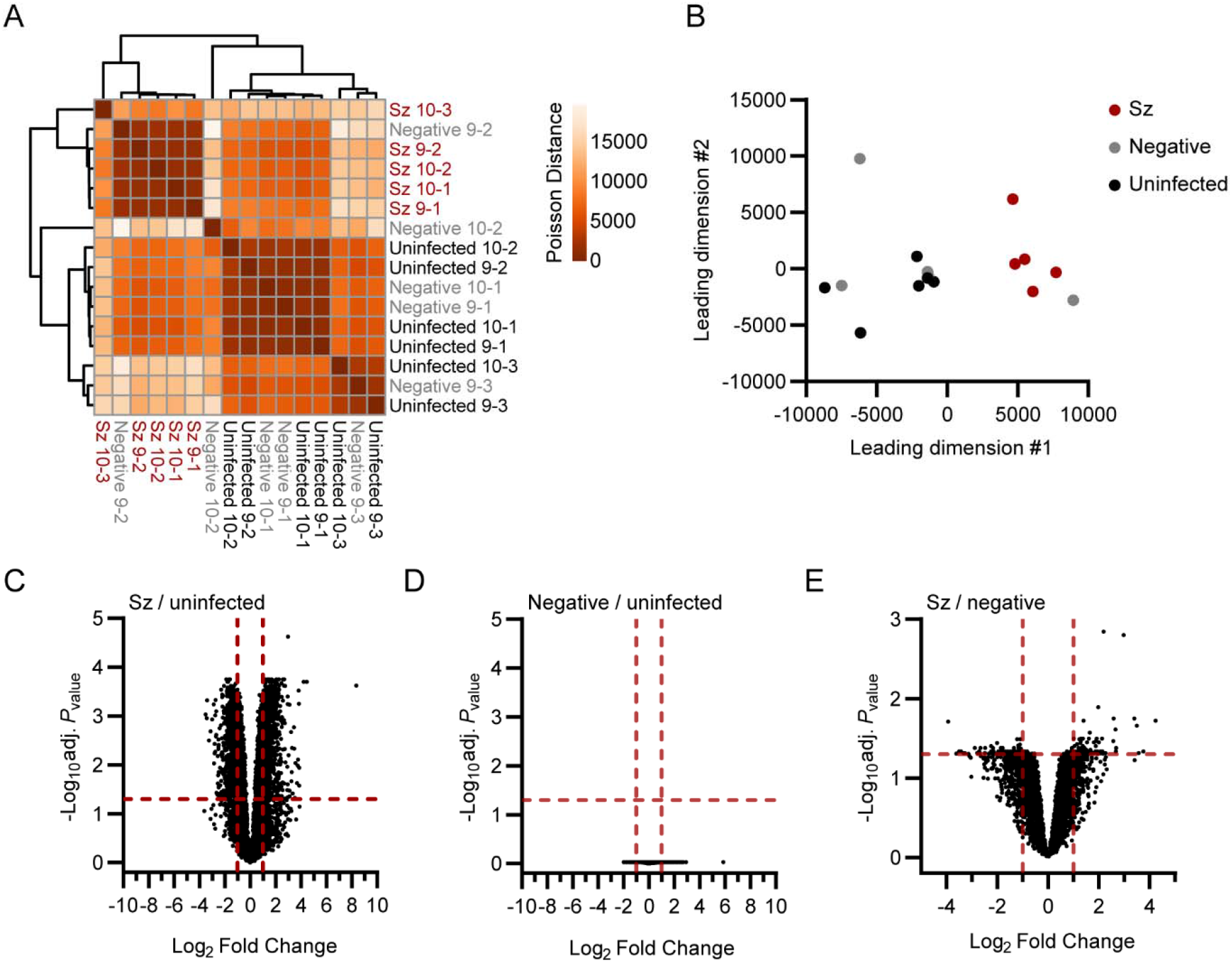
Transcriptomic analysis of primary rhesus macaque hepatocytes infected with *P. cynomolgi* schizonts. Heat map (A) and multidimensional scaling (MDS) plot (B) showing sample-to-sample separation of cells infected with schizonts (Sz, in red), uninfected bystander (Negative, in grey) cells and cells from uninfected samples (Uninfected, in black). Volcano plots showing mean log_2_ fold changes and -log_10_ adjusted *P*_values_ (adj. *P*_value_) for cells infected with schizonts in comparison to uninfected samples (C), negative cells in comparison to uninfected samples (D) and cells infected with schizonts in comparison to negative cells (E). Horizontal and vertical dotted red lines indicate adjusted *P*_values_ less than 0.05 (or -log10 (adj. *P*_value_) greater than 1.3) and absolute log_2_ fold changes greater than 1, respectively.

Thousands of genes were found to be significantly modulated (i.e., genes associated with absolute log_2_ fold change >1 and adjusted *P*_value_ < 0.05) in schizont-infected hepatocytes in comparison to uninfected samples (Figure 1C and Figure 1 – Source data 1). No statistically significant changes in gene expression were detected in uninfected bystander (negative) cells in comparison to uninfected samples (Figure 1D). However, several of the most modulated transcripts in schizont-infected cells were also modulated, although to a lesser extent and not significantly, in negative cells relative to uninfected samples (Figure 1 – Supplementary Table 1). Relatively fewer transcripts were associated with statistically significant changes in schizont-infected vs. negative cells (Figure 1E and Figure 1 – Source data 2). Interestingly, transcripts modulated in schizont-infected cells vs. uninfected samples and schizont-infected cells vs. negative cells showed some overlap for the 25 most upregulated (e.g., CXCL19, ADH7 and RGS1) and downregulated (e.g., HOXB9, RNF165 and LGALS7) genes (Figure 1 –Supplementary Table 1 and Figure 1 – Supplementary Table 2). Overall, differentially expressed genes were detected in schizont-infected hepatocytes in comparison to both uninfected samples and uninfected bystander cells, but no significant modulations were detected in uninfected bystander cells in comparison to cells from uninfected samples.

**Figure 1 – Supplementary Table 1.** Most up- and down-regulated genes in hepatocytes infected with *P. cynomolgi* schizonts in comparison to uninfected samples *^a^*

| Gene symbol | Gene description | Sz / uninfected<br>Fold change | Adj. *P*<sub>value</sub> | Negative / uninfected<br>Fold change | Adj. *P*<sub>value</sub> |
| --- | --- | --- | --- | --- | --- |
| UPREGULATED GENES |  |  |  |  |  |
| *CXCL9* | C-X-C motif chemokine ligand 9 | 327.62 | < 0.05 | 58.00 | NS |
| *ADH7* | Alcohol dehydrogenase 7 (class IV), μ or σ polypeptide | 22.10 | < 0.05 | 2.08 | NS |
| *RANBP3L* | RAN binding protein 3 like | 18.64 | < 0.05 | -1.16 | NS |
| *DOCK10* | Dedicator of cytokinesis 10 | 14.62 | < 0.05 | 3.63 | NS |
| *MAP2* | Microtubule associated protein 2 | 14.58 | < 0.05 | 2.42 | NS |
| *CXCL11* | C-X-C motif chemokine ligand 11 | 14.24 | < 0.05 | 4.38 | NS |
| *RGS1* | Regulator of G protein signaling 1 | 14.14 | < 0.05 | 2.33 | NS |
| *RGS18* | Regulator of G protein signaling 18 | 13.51 | < 0.05 | 2.65 | NS |
| *IDO1* | Indoleamine 2,3-dioxygenase 1 | 13.40 | < 0.05 | 3.89 | NS |
| *GPR65* | G protein-coupled receptor 65 | 12.89 | < 0.05 | 3.84 | NS |
| *DOCK11* | Dedicator of cytokinesis 11 | 11.91 | < 0.05 | 2.37 | NS |
| *MXRA5* | Matrix remodeling associated 5 | 11.50 | < 0.05 | 2.49 | NS |
| *TDRD6* | Tudor domain containing 6 | 11.07 | < 0.05 | 2.83 | NS |
| *DLG2* | Disks large MAGUK scaffold protein 2 | 10.79 | < 0.05 | -1.76 | NS |
| *SYTL5* | Synaptotagmin like 5 | 10.22 | < 0.05 | 3.29 | NS |
| *SLC12A5* | Solute carrier family 12 member 5 | 9.86 | < 0.05 | 1.66 | NS |
| *MRC1* | Mannose receptor C-type 1 | 9.85 | < 0.05 | 2.15 | NS |
| *CLIC2* | Chloride intracellular channel 2 | 9.70 | < 0.05 | 3.22 | NS |
| *RGS2* | Regulator of G protein signaling 2 | 9.54 | < 0.05 | 2.39 | NS |
| *MYBL1* | MYB proto-oncogene like 1 | 9.40 | < 0.05 | 3.14 | NS |
| *CCL19* | C-C motif chemokine ligand 19 | 8.61 | < 0.05 | 1.80 | NS |
| *DKK3* | Dickkopf WNT signaling pathway inhibitor 3 | 8.56 | < 0.05 | 1.27 | NS |
| *SIRPB2* | Signal regulatory protein beta 2 | 8.55 | < 0.05 | 2.73 | NS |
| *SLC17A8* | Solute carrier family 17 member 8 | 8.19 | < 0.05 | 1.10 | NS |
| *ACKR4* | Atypical chemokine receptor 4 | 7.88 | < 0.05 | -1.04 | NS |
| DOWNREGULATED GENES |  |  |  |  |  |
| *HOXB9* | Homeobox B9 | -11.47 | < 0.05 | -1.31 | NS |
| *RNF165* | Ring finger protein 165 | -10.61 | < 0.05 | -1.24 | NS |
| *LGALS7* | Galectin 7 | -10.61 | < 0.05 | -1.11 | NS |
| *C1orf210* | Chromosome 1 open reading frame 210 | -9.45 | < 0.05 | -1.20 | NS |
| *RHBG* | Rh family B glycoprotein | -8.76 | < 0.05 | -1.62 | NS |
| *RND2* | Rho family GTPase 2 | -8.01 | < 0.05 | -1.26 | NS |
| *CCDC103* | Coiled-coil domain containing 103 | -7.37 | < 0.05 | -1.12 | NS |
| *CLIP2* | CAP-Gly domain containing linker protein 2 | -7.25 | < 0.05 | -1.30 | NS |
| *NPB* | Neuropeptide B | -7.02 | < 0.05 | 1.04 | NS |
| *STAC* | SH3 and cysteine rich domain | -7.00 | < 0.05 | -2.16 | NS |
| *STRA6* | Signaling receptor and transporter of retinol STRA6 | -6.80 | < 0.05 | -1.36 | NS |
| *VILL* | Villin like | -6.75 | < 0.05 | -1.53 | NS |
| *APOE* | Apolipoprotein E | -6.66 | < 0.05 | -1.73 | NS |
| *SLC25A47* | Solute carrier family 25 member 47 | -6.49 | < 0.05 | -1.64 | NS |
| *H4C3* | H4 clustered histone 3 | -6.43 | < 0.05 | 1.15 | NS |
| *FBLN1* | Fibulin 1 | -6.36 | < 0.05 | -1.31 | NS |
| *FXYD2* | FXYD domain containing ion transport regulator 2 | -6.27 | < 0.05 | -1.85 | NS |
| *FXYD1* | FXYD domain containing ion transport regulator 1 | -5.94 | < 0.05 | -1.83 | NS |
| *SOAT2* | Sterol O-acyltransferase 2 | -5.91 | < 0.05 | -2.04 | NS |
| *FXYD-6/-2* | FXYD6-FXYD2 readthrough | -5.83 | < 0.05 | -1.68 | NS |
| *LGALS4* | Galectin 4 | -5.76 | < 0.05 | -1.48 | NS |
| *AQP7* | Aquaporin 7 | -5.73 | < 0.05 | -2.34 | NS |
| *C2CD4C* | C2 calcium dependent domain containing 4C | -5.73 | < 0.05 | -1.08 | NS |
| *CPT1B* | Carnitine palmitoyltransferase 1B | -5.69 | < 0.05 | -1.47 | NS |
| *RADIL* | Rap associating with DIL domain | -5.68 | < 0.05 | -1.24 | NS |

<sup>a</sup> Sz, schizont; Adj. *P*-values, adjusted *P*-values; NS, non-significant.

**Figure 1 – Supplementary Table 2.** Most up- and down-regulated genes in hepatocytes infected with *P. cynomolgi* schizonts in comparison to uninfected bystander (negative) cells **Figure 1 - Source data 1.** Transcriptomic analysis of schizont-infected and uninfected bystander (negative) primary rhesus macaque hepatocytes in comparison to uninfected samples. This file includes read counts, fold changes, *P* and adjusted *P* values as well as lists of genes significantly up- or down-regulated. **Figure 1 - Source data 2.** Transcriptomic analysis of primary rhesus macaque hepatocytes infected with schizonts in comparison to uninfected bystander (negative) cells. This file includes fold changes, *P* and adjusted *P*_values_ as well as lists of genes significantly up- or down-regulated.

| Gene symbol | Gene description | Fold change |
| --- | --- | --- |
| <u>UPREGULATED GENES</u> |  |  |
| *RANBP3L* | RAN binding protein 3 like | 18.95 |
| *DLG2* | Discs large MAGUK scaffold protein 2 | 13.53 |
| *LGR5* | Leucine rich repeat containing G protein-coupled receptor 5 | 11.95 |
| *SLC17A8* | Solute carrier family 17 member 8 | 11.35 |
| *ADH7* | Alcohol dehydrogenase 7 (class IV), mu or sigma polypeptide | 10.60 |
| *ACKR4* | Atypical chemokine receptor 4 | 7.92 |
| *CCL19* | C-C motif chemokine ligand 19 | 6.42 |
| *TTN* | Titin | 6.40 |
| *MAP2* | Microtubule associated protein 2 | 6.30 |
| *CCL5* | C-C motif chemokine ligand 5 | 6.00 |
| *SLC16A12* | Solute carrier family 16 member 12 | 5.99 |
| *MINAR1* | Membrane integral NOTCH2 associated receptor 1 | 5.97 |
| *ACSBG1* | Acyl-CoA synthetase bubblegum family member 1 | 4.82 |
| *RGS1* | Regulator of G protein signaling 1 | 4.64 |
| *TMEM255A* | Transmembrane protein 255A | 4.56 |
| *ADAM21* | ADAM metalloproteinase domain 21 | 4.49 |
| *PLCE1* | Phospholipase C epsilon 1 | 4.40 |
| *GCLC* | Glutamate-cysteine ligase catalytic subunit | 4.37 |
| *LRRN3* | Leucine rich repeat neuronal 3 | 4.22 |
| *FGF7* | Fibroblast growth factor 7 | 4.20 |
| *FAM171B* | Family with sequence similarity 171 member B | 4.19 |
| *GLI1* | GLI family zinc finger 1 | 4.16 |
| *ADGRF5* | Adhesion G protein-coupled receptor F5 | 4.15 |
| *HRH4* | Histamine receptor H4 | 4.11 |
| *MEF2C* | Myocyte enhancer factor 2C | 4.03 |
| <u>DOWNREGULATED GENES</u> |  |  |
| *HOXB9* | Homeobox B9 | -15.21 |
| *H1-1* | H1.1 linker histone, cluster member | -12.22 |
| *NPB* | Neuropeptide B | -11.27 |
| *LGALS7* | Galectin 7 | -11.22 |
| *C1orf210* | Chromosome 1 open reading frame 210 | -10.77 |
| *OPRPN* | Opiorphin prepropeptide | -10.07 |
| *KLK10* | Kallikrein related peptidase 10 | -9.69 |
| *CCDC71* | Coiled-coil domain containing 71 | -8.73 |
| *WDR86* | WD repeat domain 86 | -7.39 |
| *H4C3* | H4 clustered histone 3 | -6.87 |
| *PRG2* | Proteoglycan 2, pro-eosinophil major basic protein | -6.54 |
| *RNF165* | Ring finger protein 165 | -6.43 |
| *GRAMD2A* | GRAM domain containing 2A | -6.30 |
| *H1-4* | H1.4 linker histone, cluster member | -5.61 |
| *TRIM15* | Tripartite motif containing 15 | -5.54 |
| *STAR* | Steroidogenic acute regulatory protein | -5.37 |
| *C1QTNF2* | C1q and TNF related 2 | -5.03 |
| *TMIE* | Transmembrane inner ear | -4.86 |
| *CCDC103* | Coiled-coil domain containing 103 | -4.74 |
| *STRA6* | Signaling receptor and transporter of retinol STRA6 | -4.71 |
| *CDKN2A* | Cyclin dependent kinase inhibitor 2A | -4.59 |
| *GNAT2* | G protein subunit alpha transducin 2 | -4.59 |
| *RASL11A* | RAS like family 11 member A | -4.50 |
| *MMP7* | Matrix metalloproteinase 7 | -4.48 |
| *SLPI* | Secretory leukocyte peptidase inhibitor | -4.46 |

### Pathway analysis of differentially expressed genes in schizont-infected hepatocytes in comparison to cells from uninfected samples

To focus on the most robustly modulated genes, filtered-by-threshold pathway enrichment analyses were performed using genes associated with statistically significant changes in schizont-infected cells versus uninfected samples (Figure 1C and Figure 1 – Source data 1) and Metascape (Zhou et al., 2019). The 20 most enriched Metascape ontology clusters in genes upregulated by schizont-infected cells (Figure 2 – Source data 1) are represented in a histogram (Figure 2A) and in a network layout that connects enriched terms with a certain level of similarity (Figure 2B). Interestingly, 7 of the most significantly enriched ontology clusters were interconnected and shared genes associated with the response to DNA damage, cell division, microtubule-based process, regulation of cell cycle, MTOC organization, regulation of cytoskeleton organization and regulation of organelle organization. Two other ontology clusters were also interconnected and included genes involved in regulation of the cellular stress response and in regulation of protein kinase activity. The remaining most significantly enriched ontology clusters were discrete and related to other pathways such as RHO GTPase cycle, response to *Herpes simplex* virus 1 (HSV-1) infection, proteolysis involved in catabolism, membrane trafficking and response to cytokines. Ten ontology clusters were further selected and the expression level of the 25 most up-regulated genes were visualized for each of these clusters in both schizont-infected and uninfected bystander (negative) cells in comparison to uninfected cells (Figures 2C-2L). While most groups of genes were more strongly and homogenously upregulated in schizont-infected cells, several genes were also expressed at noticeable levels in negative cells (e.g., CLSPN, PDCD1LG2 and CXCL9). The results thus suggested that schizont-infected hepatocytes upregulate multiple biological pathways in comparison to cells from uninfected samples and, in most cases but not always exclusively, in comparison to uninfected bystander cells.

**Figure 2.**
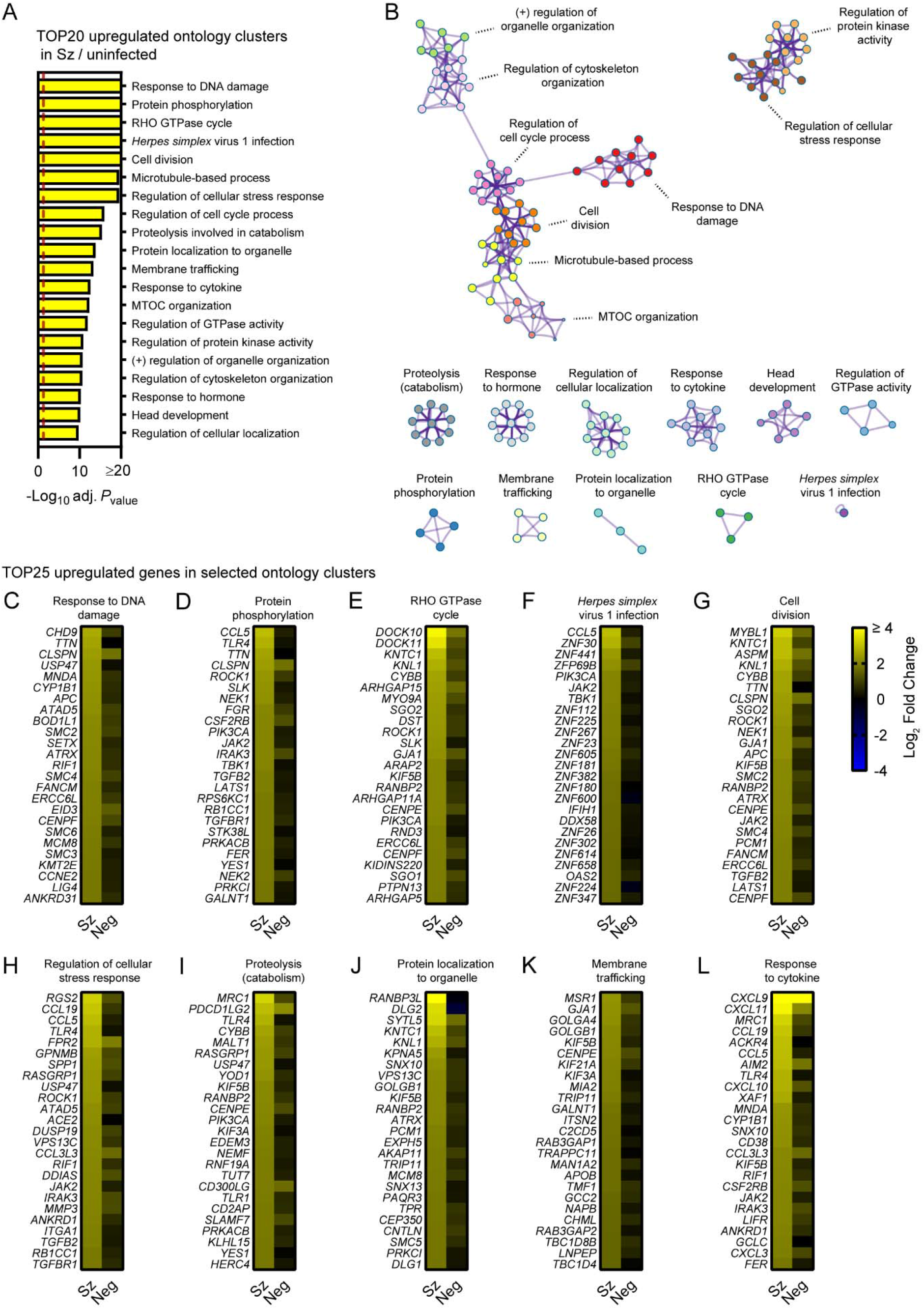
Pathway analysis of genes significantly upregulated in primary hepatocytes infected with *P. cynomolgi* schizonts in comparison to cells from uninfected samples. (A) Most enriched ontology clusters for genes significantly upregulated in primary rhesus macaque hepatocytes infected with schizonts. The vertical dotted red line indicates adjusted *P*_values_ less than 0.05 (or -log10 (adj. *P*_value_) greater than 1.3). The most significantly enriched term of each ontology clusters was selected as the cluster label. (B) Subset of representative terms were selected to generate a network of enriched ontology clusters. Terms are represented by circle nodes with colors that shows belonging to a specific cluster. Circles nodes with similarity scores > 0.3 are linked with edges. (C-L) Heat maps of log_2_ fold changes for the 25 most upregulated genes for selected ontology clusters. Data are shown for schizont-infected and uninfected bystander (negative) cells in comparison to uninfected samples. Some annotation terms are abbreviated or modified for a purpose of presentation. **Figure 2. Source data 1.** Term annotation and pathway enrichment analysis for genes significantly upregulated in primary rhesus macaque hepatocytes infected with schizonts in comparison to uninfected samples. This analysis was performed with Metascape.

Metascape pathway analysis of genes significantly downregulated in schizont-infected cells vs. uninfected samples showed the enrichment of two networks of ontology clusters that relate to protein translation and ribosomes, and metabolic pathways and mitochondria, respectively (Figures 3A and 3B, Figure 3 – Source data 1). Three other ontology clusters were distinct and included genes associated with γ-carboxylation, hypusine formation and arylsulfatase activation, regulation of peptidase activity and regulation of growth (Figure 3B). Interestingly, Metascape analyses also revealed that genes associated with the specific transcriptional signature of the liver tissue (PaGenBase (Pan et al., 2013)) were significantly enriched among transcripts downregulated in schizont-infected cells (Figure 3 – Figure supplement 1). Analysis of the 25 most down-regulated genes for selected ontology clusters showed robust and overall consistent gene repression in schizont-infected cells in comparison to both uninfected samples and uninfected bystander cells (Figures 3C-3L). These results suggested that schizont-infected hepatocytes significantly downregulate genes involved in multiple biological pathways, including pathways associated with protein translation and metabolic processes, as well as genes associated with the transcriptional signature of the liver tissue.

**Figure 3.**
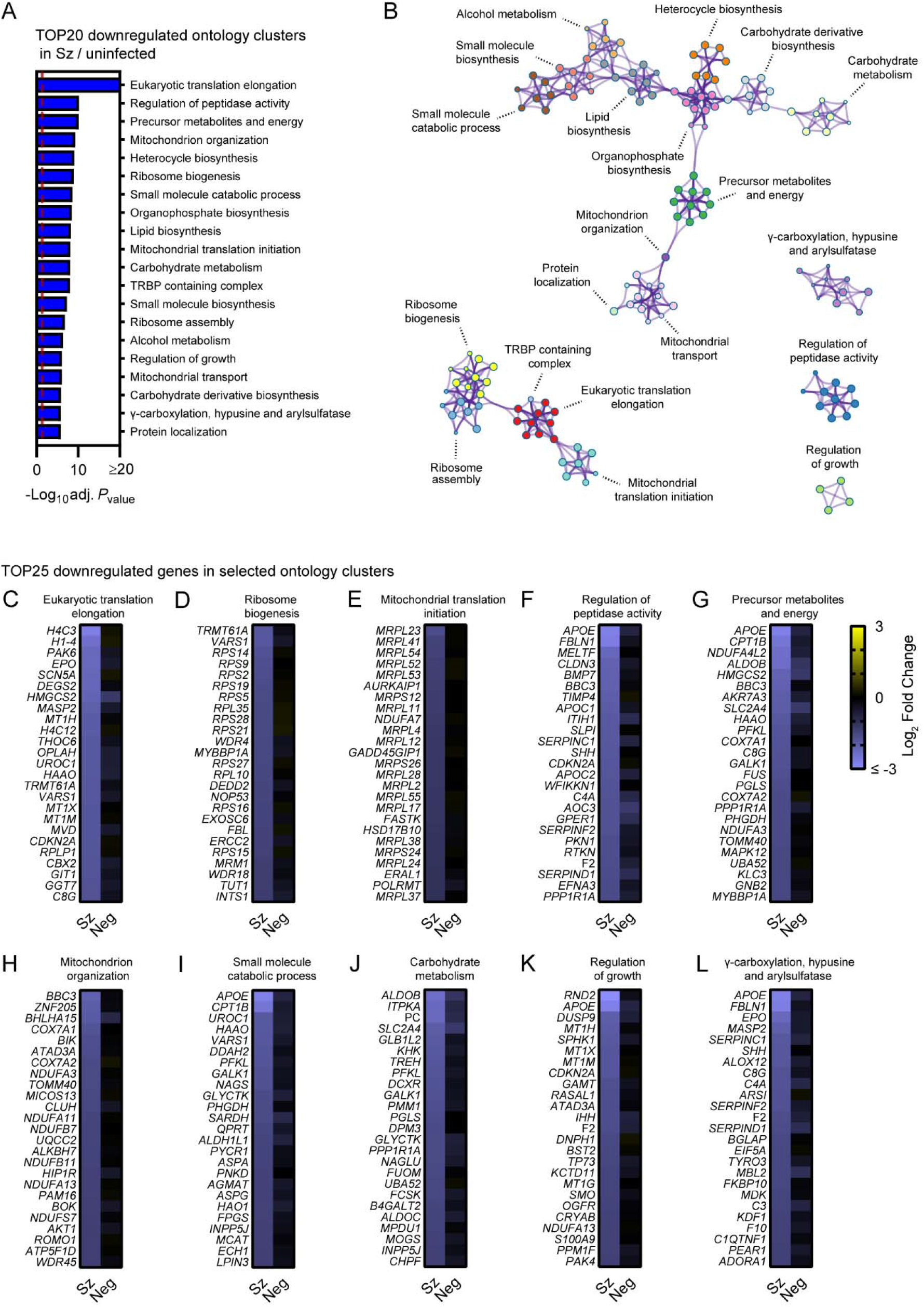
Pathway analysis of genes significantly downregulated in primary hepatocytes infected with *P. cynomolgi* schizonts in comparison to cells from uninfected samples. (A) Most enriched ontology clusters for genes significantly downregulated in primary rhesus macaque hepatocytes infected with schizonts. The vertical dotted red line indicates adjusted *P*_values_ less than 0.05 (or -log10 (adj. *P*_value_) greater than 1.3). The most significantly enriched term of each ontology clusters was selected as the cluster label. (B) Subset of representative terms were selected to generate a network of enriched ontology clusters. Terms are represented by circle nodes with colors that shows belonging to a specific cluster. Circles nodes with similarity scores > 0.3 are linked with edges. (C-L) Heat maps of log_2_ fold changes for the 25 most downregulated genes of selected ontology clusters. Data are shown for schizont-infected and uninfected bystander (negative) cells in comparison to uninfected samples. Some annotation terms are abbreviated or modified for a purpose of presentation. **Figure 3. Source data 1.** Term annotation and pathway enrichment analysis for genes significantly downregulated in primary rhesus macaque hepatocytes infected with schizonts in comparison to uninfected samples. This analysis was performed with Metascape.

**Figure 3 – Figure supplement 1.**
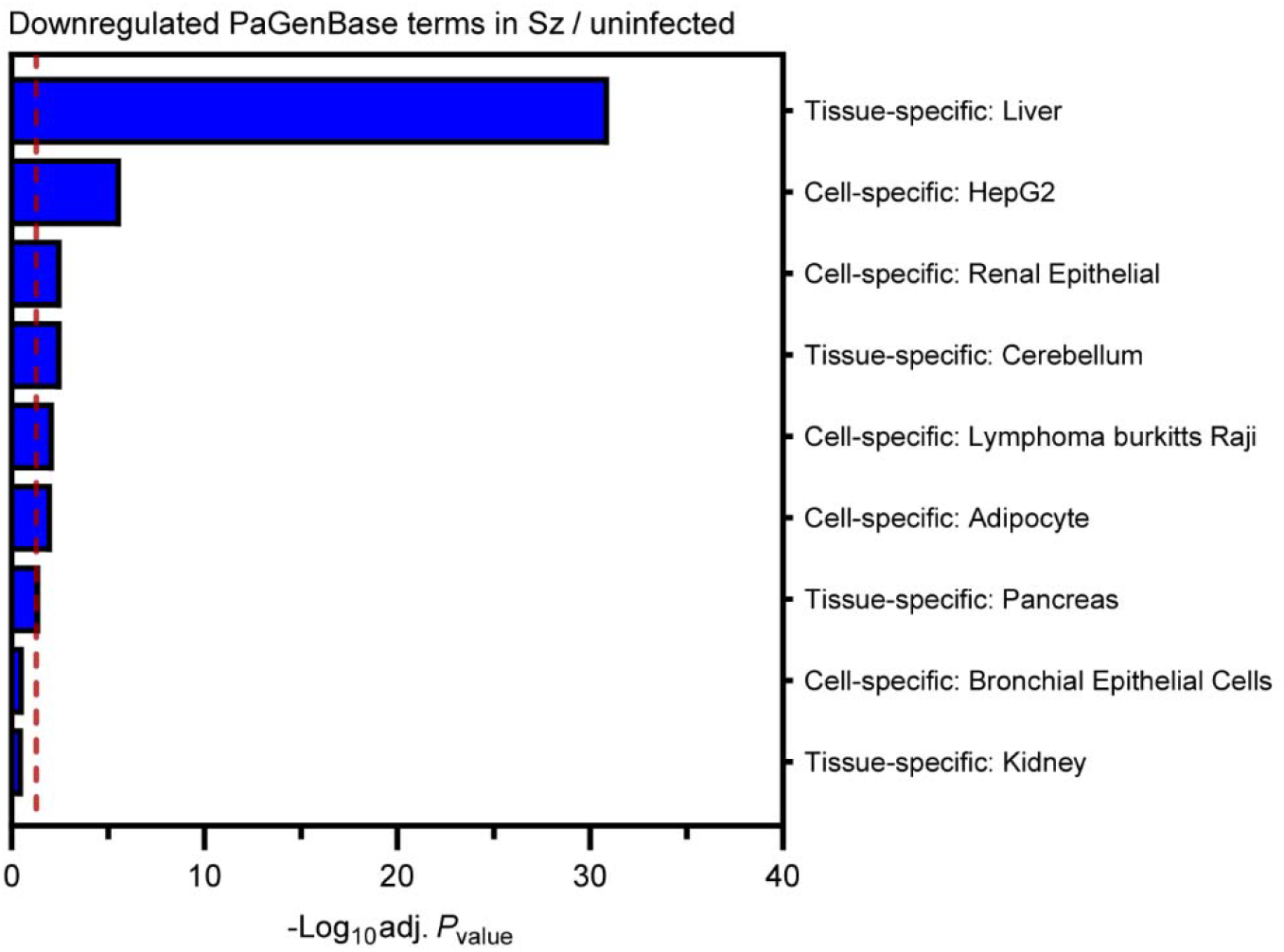
Most enriched PaGenBase (Pattern Gene Database) terms for genes significantly downregulated in hepatocytes infected with schizonts in comparison to uninfected samples. The vertical dotted red line indicates adjusted *P*_values_ less than 0.05 (or -log10 (adj. *P*_value_) greater than 1.3).

### Pathway analysis of differentially expressed genes in schizont-infected hepatocytes in comparison to uninfected bystander cells

To confirm results from the previous section as well as to identify transcriptional changes strictly modulated in infected cells, Metascape pathway analysis was performed for the transcripts significantly modulated in schizont-infected vs. uninfected bystander (negative) cells. This analysis revealed enrichment of ontology clusters associated with specific biological processes in both upregulated (e.g., RHO GTPase cycle and protein phosphorylation) (Figures 4A and Figure 4 – Source data 1) and downregulated (e.g., ribosome and oxidative phosphorylation) genes (Figure 4B and Figure 4 – Source data 1). This analysis also confirmed the downregulation of genes associated with the liver-specific transcriptional signature in *P. cynomolgi*-infected hepatocytes (Figure 4 – Figure supplement 1).

**Figure 4.**
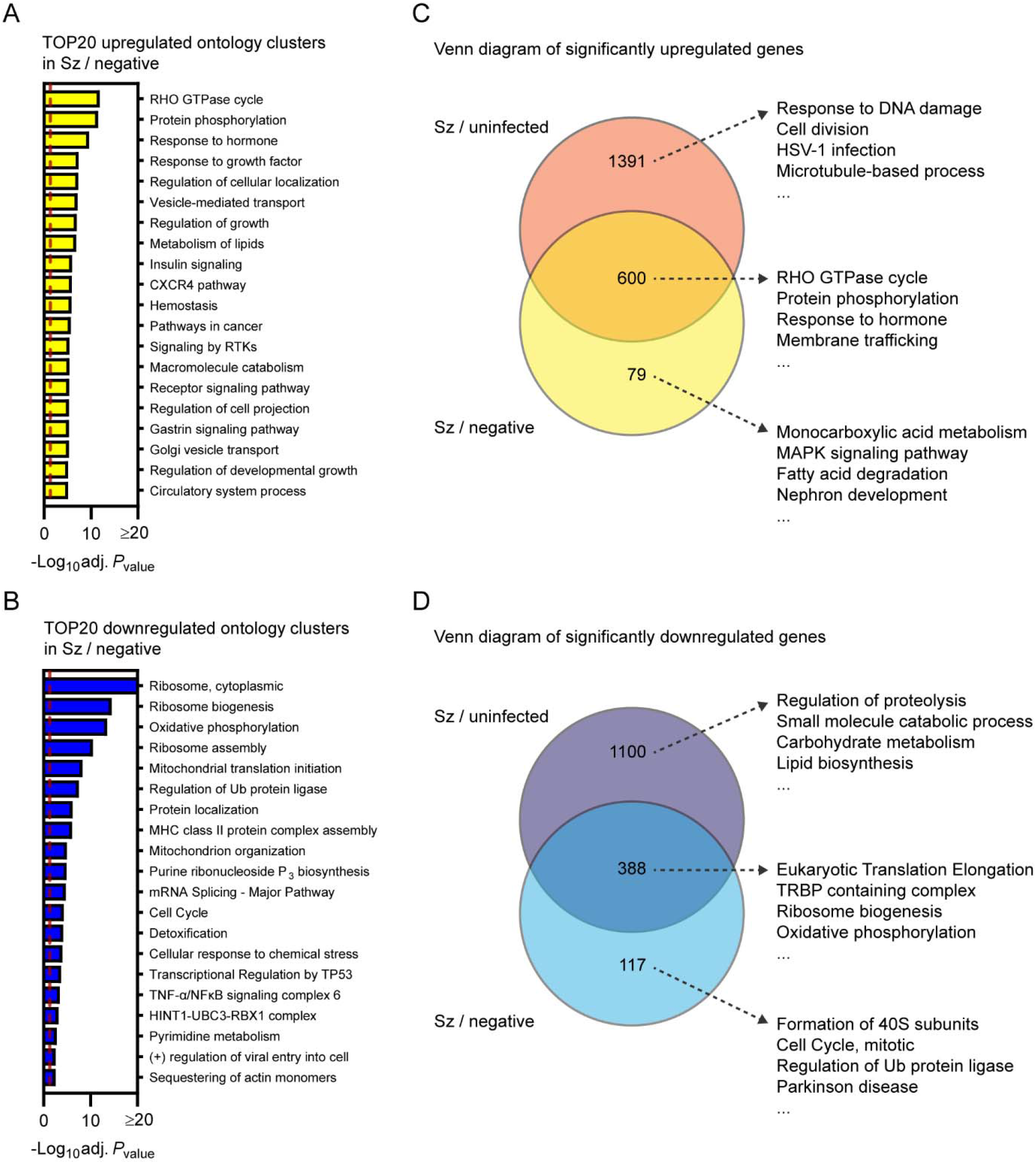
Transcriptional response of primary hepatocytes infected with *P. cynomolgi* schizonts in comparison to uninfected bystander cells. Most enriched ontology clusters for genes significantly upregulated (A) and downregulated (B) in hepatocytes infected with schizonts in comparison to uninfected bystander (negative) cells. Venn diagram analyses comparing upregulated (C) and downregulated (D) genes in schizont-infected hepatocytes vs. uninfected samples or vs. negative cells. The 4 most enriched ontology clusters are shown for each subgroup of genes. Dotted red lines indicate adjusted *P*_values_ less than 0.05 (or -log10 (adj. *P*_value_) greater than 1.3). Some annotation terms are abbreviated or modified for a purpose of presentation. **Figure 4 – Source data 1.** Term annotation and pathway enrichment analysis for genes significantly modulated in primary rhesus macaque hepatocytes infected with schizonts in comparison to uninfected bystander (negative) cells. **Figure 4 – Source data 2.** Venn diagram analysis lists, term annotation and pathway enrichment analysis for significantly upregulated genes in hepatocytes infected with schizonts in comparison to uninfected samples or uninfected bystander (negative) cells. **Figure 4 – Source data 3.** Venn diagram analysis lists, term annotation and pathway enrichment analysis for significantly downregulated genes in hepatocytes infected with schizonts in comparison to uninfected samples or uninfected bystander (negative) cells. **Figure 4 – Source data 4.** Gene Set Enrichment Analysis (GSEA) for the transcriptional profiling of schizont-infected cells vs. uninfected samples, uninfected bystander (negative) cells vs. uninfected samples and schizont-infected cells vs. negative cells. This file includes lists for all and selected gene sets (i.e., Gene Ontology (GO), the KEGG pathway and the Reactome pathway databases) from the MSigDB collections. Only datasets associated with a False Discovery Rate (FDR) < 0.05 are shown.

**Figure 4 – Figure supplement 1.**
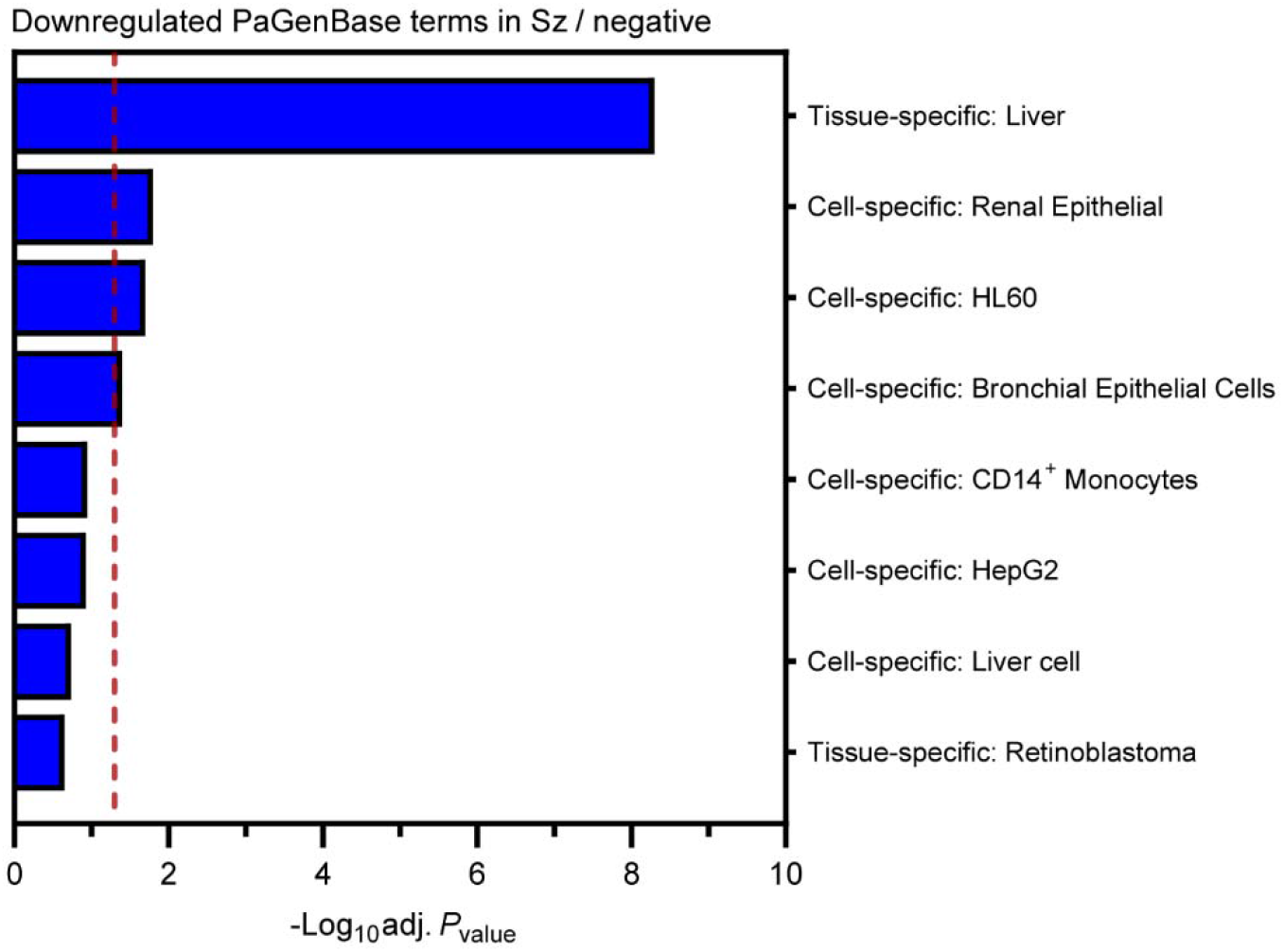
Most enriched PaGenBase (Pattern Gene Database) terms for genes significantly downregulated in hepatocytes infected with schizonts in comparison to uninfected bystander (negative) cells. The vertical dotted red line indicates adjusted *P*_values_ less than 0.05 (or -log10 (adj. *P*_value_) greater than 1.3).

Venn diagram analysis was used to compare the differentially expressed genes in schizont-infected hepatocytes vs. uninfected samples and in schizont-infected hepatocytes vs. negative cells, and the genes of each subgroups were then submitted to a Metascape pathway analysis (Figures 4C and 4D). Genes only upregulated in schizont-infected cells vs. uninfected samples were associated with specific ontology clusters, such as response to DNA damage, cell division, HSV-1 infection, and microtubule-based process (Figures 4C and FIG. 4 – Source data 2) and might either represent genes that are also induced in negative cells or more sensitive to noise (i.e., and more impacted by the greater variability between negative samples in comparison to uninfected samples). Genes upregulated in both schizont-infected cells vs. uninfected samples and schizont-infected cells vs. negative cells were associated with other ontology clusters, such as Rho GTPase cycle, protein phosphorylation, response to hormone and membrane trafficking (Figure 4C and Figure 4 – Source data 2), and represent transcripts specifically induced in infected cells. Results similarly show the specific enrichment of ontology clusters for genes only downregulated in schizont-infected vs. uninfected samples (e.g., regulation of proteolysis and small molecule catabolic process) or downregulated in both schizont-infected vs. uninfected samples and schizont-infected vs. negative cells (e.g., eukaryotic translation elongation, TRBP containing complex, ribosome biogenesis and oxidative phosphorylation) (Figure 4D and Figure 4 – Source data 3). Ontology clusters enriched in genes specifically modulated by schizont-infected cells vs. negative cells were more difficult to interpret but might reflect changes indirectly induced by the infection in the uninfected bystander cell population (Figures 4C and 4D). Altogether, these results showed that changes in different biological processes are detected in schizont-infected hepatocytes depending on whether the data are compared to uninfected samples or to uninfected bystander cells.

It was hypothesized that differences in the transcriptional profiles of samples normalized to either uninfected samples or negative cells were caused by the modulation of specific transcripts, although below our previous threshold for statistical significance (absolute log_2_ fold change >1 and adjusted *P*_value_ < 0.05), in the uninfected bystander (negative) cell population. Accordingly, numerous transcripts were differentially expressed in negative cells in comparison to uninfected samples once the data were filtered using a more permissive threshold (e.g., absolute log_2_ fold change >1 and *P*_value_ < 0.05) (Figure 4 – Figure supplement 2). To better understand the transcriptional changes in negative cells, data were analyzed using Gene Set Enrichment Analysis (GSEA) (Figure 4 – Source data 4), a threshold-free method that does not require prior filtering of genes based on statistically significant transcriptional changes (Liberzon et al., 2015; Liberzon et al., 2011; Subramanian et al., 2005). Comparison of GSEA pathways enriched in schizont-infected cells and negative cells using all gene sets from the MSigDB collection showed a considerable overlap for both the upregulated and, to a lesser extent, downregulated pathways (Figure 4 – Figure supplement 3). Interestingly, among selected gene sets (i.e., Gene Ontology (GO), the KEGG pathway and the Reactome pathway databases), GSEA pathways that relate to chromosome biology were significantly enriched in both schizont-infected and negative cells in comparison to uninfected samples, but not in schizont-infected vs. negative cells (Figure 4 – Figure supplement 4), demonstrating a common regulation of these pathways in infected and uninfected bystander cells. Overall, these results strongly suggested that a group of specific host processes are modulated at the transcriptional level in the uninfected bystander cell population during the liver stage of malaria infection.

**Figure 4 – Figure supplement 2.**
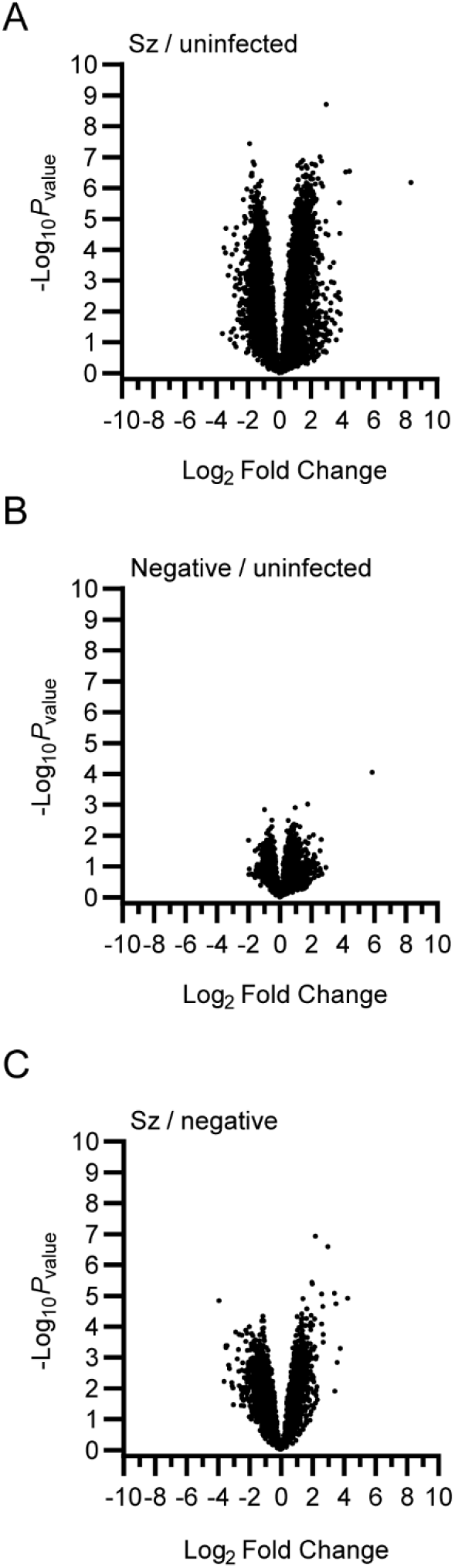
Volcano plots showing mean log_2_ fold changes and -log_10_ *P*_values_ for schizont-infected vs. uninfected samples (A), uninfected bystander (negative) cells vs uninfected samples (B) and schizont-infected vs. negative cells (C).

**Figure 4 – Figure supplement 3.**
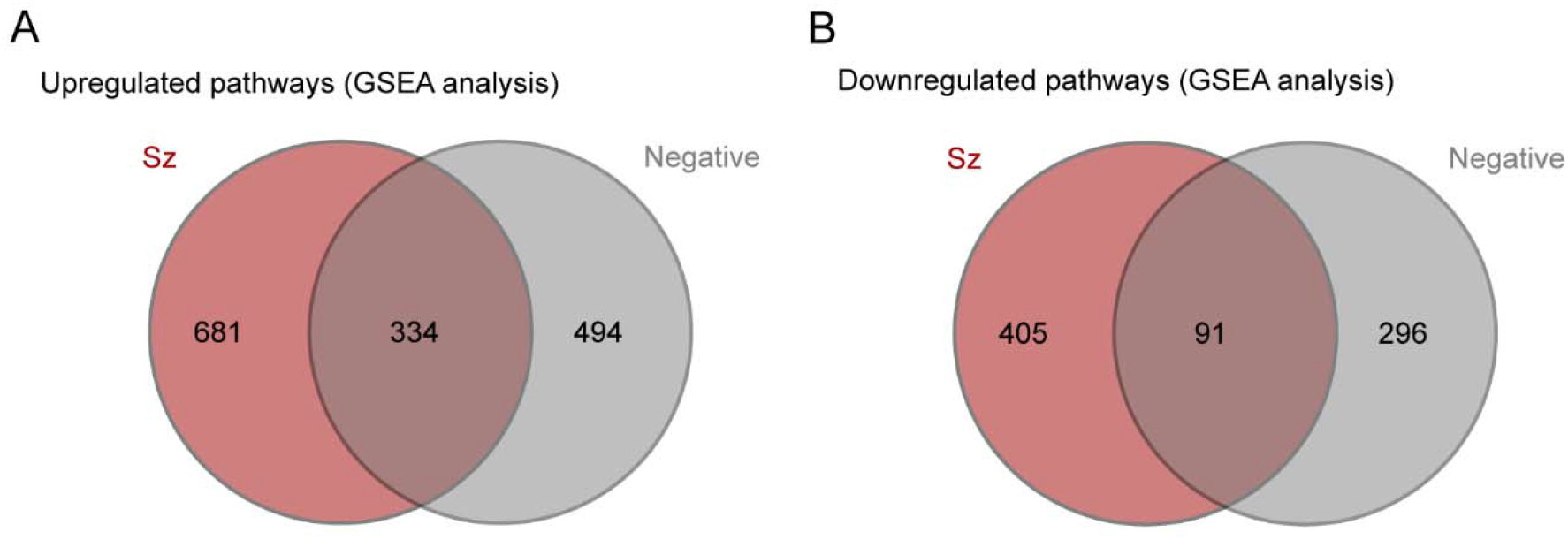
Venn diagram analyses comparing upregulated (A) and downregulated (B) gene sets / pathways from GSEA analyses in schizont-infected vs. uninfected samples and uninfected bystander (negative) cells vs. uninfected samples. All gene sets from the MSigDB collections were considered for these analyses.

**Figure 4 – Figure supplement 4.**
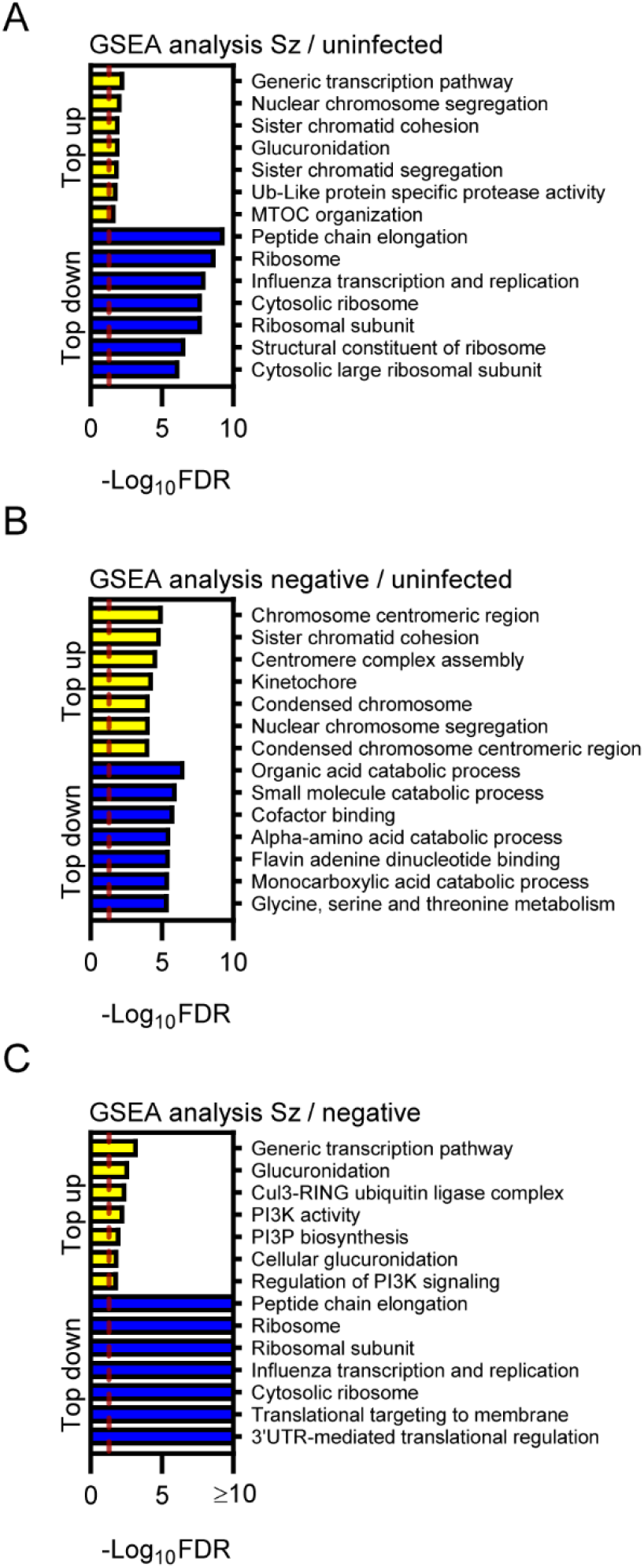
Most significantly enriched gene sets from GSEAs for the upregulated (yellow) and downregulated (blue) transcripts in schizont-infected vs. uninfected samples (A), uninfected bystander (negative) cells vs. uninfected samples (B) and schizont-infected vs. negative cells (C). Only gene sets from the Gene Ontology (GO), the KEGG pathway and the Reactome pathway databases were considered. Some annotation terms are abbreviated or modified for a purpose of presentation.

### Host transcriptional profiling of the liver stage of *Plasmodium berghei*

A dual RNA sequencing study for the liver stage of the non-relapsing rodent malaria parasite *P. berghei* was previously published (LaMonte et al., 2019). To compare our data more directly to this study, the combined host gene expression dataset for hepatocyte-like cell lines (Huh7.5.1, HC04 and HepG2) infected for 48 hours with *P. berghei* (See Supplementary Data 5 published by LaMonte et al. (2019)) was reanalyzed with the filtered-by-threshold approach used in this study (i.e., absolute log_2_ fold change >1 and adjusted *P*_value_ < 0.05) and Metascape. Similarly to *P. cynomolgi*-infected hepatocytes, cells infected with *P. berghei* upregulated genes associated with *Herpes simplex* virus 1 (HSV-1) infection, microtubule-based process and RHO GTPase cycle (Supplementary figure 1A) while downregulating genes associated with pathways such as translation, oxidative phosphorylation and metabolic processes (Supplementary figure 1B). In addition, genes associated with the specific transcriptional signature of the liver tissue (PaGenBase database) were also significantly enriched among transcripts downregulated in *P. berghei*-infected cells (Supplementary figure 2). Thus, the transcriptomes of cells infected with *P. cynomolgi* and *P. berghei* liver stage schizonts overlap and show modulation of similar biological processes.

**Supplementary figure 1.**
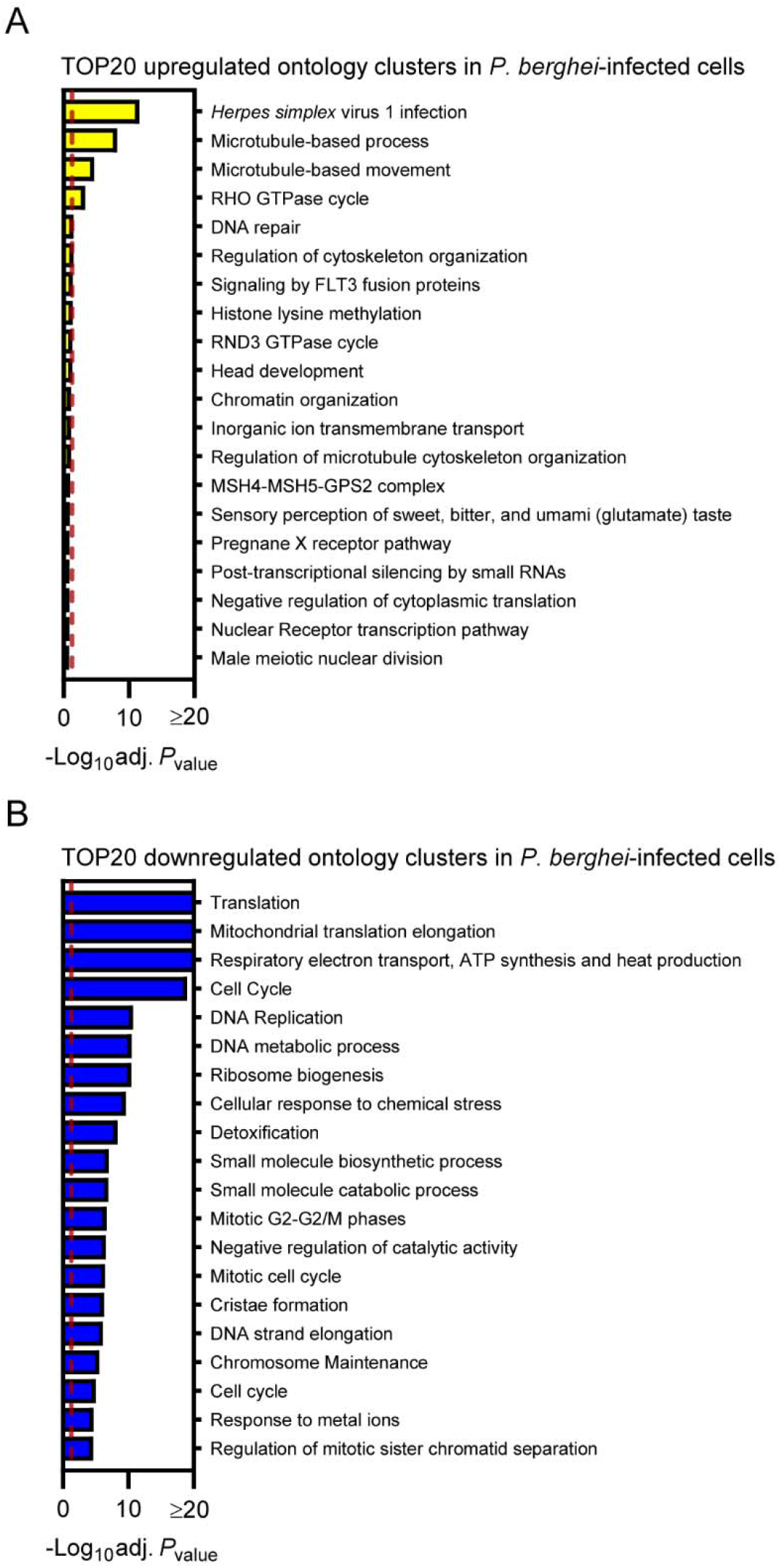
Re-analysis of the host response to *P. berghei* liver stage schizonts. Most enriched ontology clusters for genes significantly upregulated (A) and downregulated (B) in cells infected with *P. berghei* liver stage schizonts. Dotted red lines indicate adjusted *P*_values_ less than 0.05 (or -log10 (adj. *P*_value_) greater than 1.3). Some annotation terms are abbreviated or modified for a purpose of presentation.

## DISCUSSION

This study provides a snapshot of the transcriptional profile of primary rhesus macaque hepatocytes infected with the replicative form of the malaria parasite *P. cynomolgi*. It was found that schizont-infected cells modulate the expression of genes associated with multiple host processes in comparison to uninfected samples but also to uninfected bystander cells. The reason why some gene modulations were only observed in schizont-infected hepatocytes when normalized to uninfected samples but not to uninfected bystander cells is not known, and might be of technical (e.g., greater variability within the uninfected bystander cell population) or of biological (e.g., heterogeneous induction of host genes in non-infected cells) origin. However, while the comparison to uninfected samples provides a more exhaustive picture of the host response to infection, the differentially expressed genes identified from the comparison to uninfected bystander cells are more likely to be specific to schizont-infected hepatocytes.

The host pathways modulated in cells infected with the liver stages of *P. cynomolgi* and *P. berghei* considerably overlap. Similarly to *P. cynomolgi*-infected hepatocytes, our re-analysis of the dual RNAseq dataset previously published by LaMonte et al. (2019) showed that cells infected with the liver stage of *P. berghei* upregulate genes associated with *Herpes simplex* virus 1 (HSV-1) infection, microtubule-based processes and the RHO GTPase cycle (Supplementary figure 1A), and down-regulate genes associated with translation, oxidative phosphorylation and liver-specific functions (Supplementary figures 1B and 2). The upregulation of genes associated with HSV-1 infection and the response to cytokines in hepatocytes infected with *P. cynomolgi* schizonts (Figure 2) also agrees with a previous study suggesting that *P. berghei* activates innate immunity during the liver stage of infection (Liehl et al., 2014). However, as strongly suggested by another study profiling the host transcriptome during the liver stage of *P. berghei* (Posfai et al., 2018), the response to infection is not only dynamic and evolved as a function of time but is also highly dependent on the background of host cells being infected. Consequently, differences between the response of primary rhesus macaque hepatocytes and immortalized hepatocyte-like human cells to malaria parasites might be exacerbated by their respective proliferative state as well as by species-specific variations. The host pathways that are modulated independently of the host and parasite species might represent the specific transcriptional signature of schizont-infected hepatocytes and are likely to include host processes that are essential for liver stage development.

**Supplementary figure 2.**
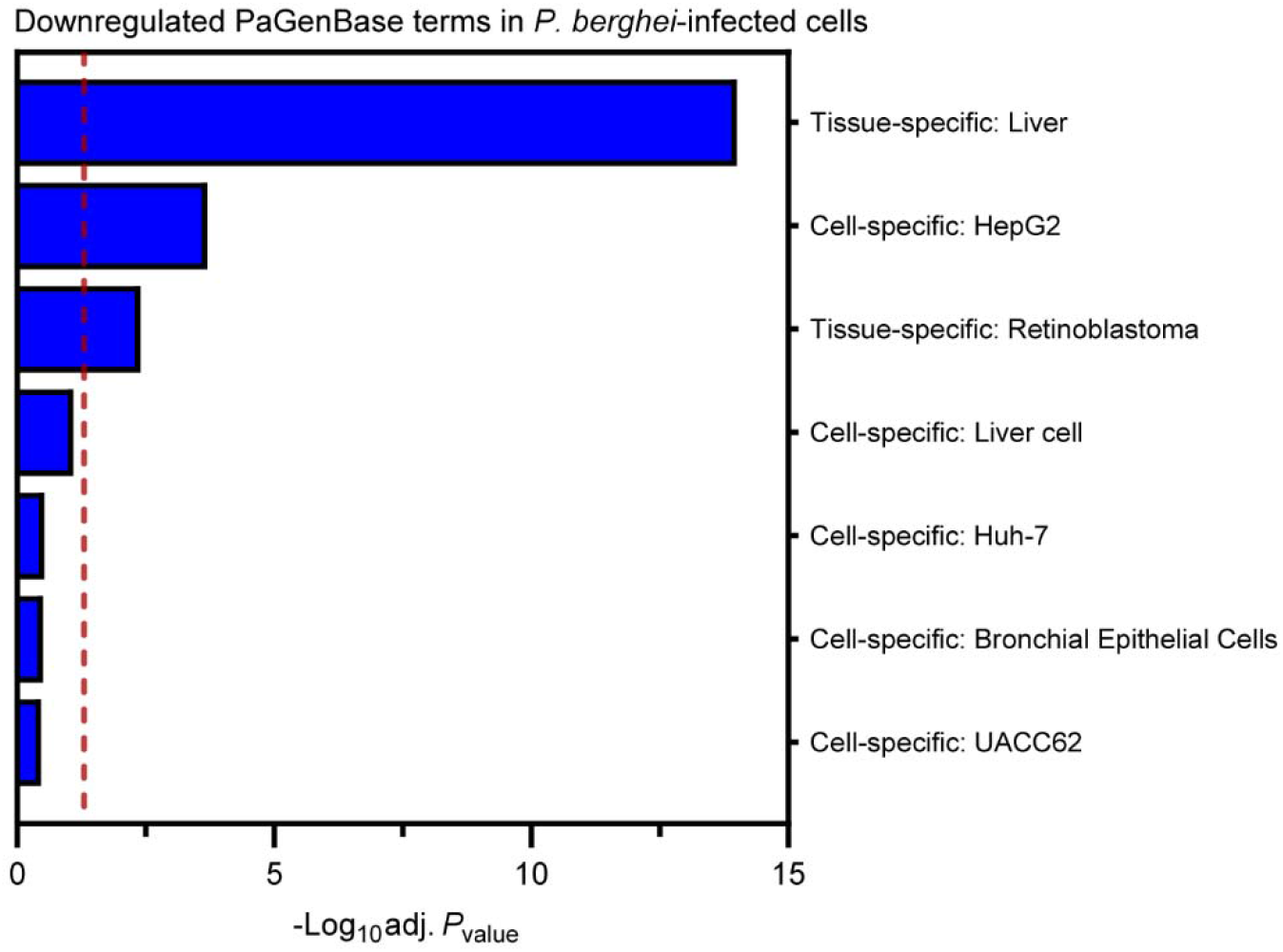
Re-analysis of the host response to *P. berghei* liver stage schizonts. Most enriched PaGenBase (Pattern Gene Database) terms for genes significantly downregulated in cells infected with *P. berghei* liver stage schizonts. The vertical dotted red line indicates adjusted *P*_values_ less than 0.05 (or -log10 (adj. *P*_value_) greater than 1.3).

Recent studies reported the host transcriptional profile of human hepatocytes infected with *P. vivax* (Mancio-Silva et al., 2022; Ruberto et al., 2022). Interestingly, Mancio-Silva et al. (2022) found that the late phase of the *P. vivax* liver stage is associated with the downregulation of liver-specific functions as well as translation, as observed for *P. cynomolgi*- and *P. berghei*-infected hepatocytes (Figures 3 and 4 and Supplementary figures 1 and 2). However, few significantly upregulated genes were observed for *P. vivax*-infected hepatocytes (Mancio-Silva et al., 2022), contrasting with the potent induction of numerous host transcripts during the liver stages of *P. berghei* (LaMonte et al., 2019) and *P. cynomolgi* (Figure 1). Moreover, the study by Ruberto et al. (2022) reported opposite modulations of host pathways (e.g., upregulations of host genes associated with translation and oxidative phosphorylation) in hepatocytes infected with *P. vivax*, which is in apparent disagreement with our results, as well as with results from others (LaMonte et al., 2019; Mancio-Silva et al., 2022). Thus, beyond the impact of malaria- and host-species specific variations, current results suggest that the hepatocyte response to infection is highly dynamic or influenced by technical aspects (e.g., the source of human hepatocytes). Accordingly, more thorough studies are required to accurately compare the transcriptome of hepatocytes infected by different species of malaria.

The lack of data for the transcriptional profiling of hepatocytes infected with *P. cynomolgi* hypnozoites is a major limitation of this study. This limitation was caused by the technical challenges of sorting hypnozoite-infected hepatocytes at later infection timepoints using the FACS-based approach (Bertschi et al., 2018). This study also only provides a snapshot of the transcriptional signature of schizont-infected cells. As such, future transcriptomic studies should be performed at different timepoints to evaluate the dynamic aspect of the host response as well as to compare the host transcriptional signature of hepatocytes infected with *P. cynomolgi* schizonts and hypnozoites. Moreover, this study did not identify any significantly modulated genes in the bystander uninfected cell population (Figure 1), which is likely caused by a heterogeneity in the transcriptome of these cells and could be better resolved using spatial transcriptomic technologies (Rao et al., 2021). Despite its limitations, this pioneering study characterized the transcriptional signature of rhesus macaque hepatocytes infected with the replicative form of *P. cynomolgi* and found similitudes with the host response induced by the liver stages of other malaria species. This study thus further validates *P. cynomolgi* as a model organism to study relapsing malaria and provides a framework to build on future research that aims at understanding host-pathogen interactions during the liver stages of malaria infection.

## MATERIALS AND METHODS

### Ethics statement

Nonhuman primates were used because no other models (in vitro or in vivo) were suitable for the aims of this project. The research protocol was approved by the local independent ethical committee conform Dutch law (BPRC Dier Experimenten Commissie, DEC, agreement number #708). Details were previously described by Voorberg-van der Wel et al. (2017).

### Preparation of samples

Cells from uninfected samples and uninfected bystander (GFP-negative, negative) cells were isolated and processed for RNA sequencing in-parallel with previously published *P. cynomolgi*-infected samples (Bertschi et al., 2018). Cells from uninfected samples were also FACS-sorted. Details were previously described (Bertschi et al., 2018; Voorberg-van der Wel et al., 2017).

### Data availability

The raw RNA-sequencing reads for infected samples analyzed in this study were previously published (Bertschi et al., 2018) and made available in the NCBI Short Read Archive (https://www.ncbi.nlm.nih.gov/sra) under accession number SRP096160. The raw RNA-sequencing reads for uninfected samples and uninfected bystander (negative) cells will be deposited into the NCBI Short Read Archive.

### Processing and normalization of RNA sequencing data

The 76-bp paired-end reads were aligned to the reference rhesus monkey genome rheMac7 (Zimin et al., 2014) and used for gene expression quantification with the Exon Quantification Pipeline (EQP) (Schuierer & Roma, 2016), as described by Voorberg-van der Wel et al. (2017). The reference rhesus monkey genome rheMac7 is annotated with human gene identification numbers. Gene raw counts were log transformed, normalized and a differential gene expression analysis was performed using the limma/voom Bioconductor pipeline (Ritchie et al., 2015) (EdgeR version 3.28.1, limma version 3.42.2) and R (version 3.6.0, 2019-04-26) using a x86_64-pc-linux-gnu (64-bit) platform running under a CentOS Linux 7 (Core).

### Sample distances, clustering and multidimensional scaling analyses

Sample distances and clustering were calculated using the Poisson distance (Love et al., 2015), the gene raw counts and RStudio (version 1.1.456). Poisson distances were also used to generate multidimensional scaling (MDS) plots on RStudio, as described by Love et al. (2015).

### Pathway enrichment and Venn diagram analyses

Pathway enrichment analyses were performed by filtering the differentially expressed genes using a threshold for statistical significance (absolute log_2_ fold change >1 and adjusted *P*_value_ < 0.05) and Metascape (Zhou et al., 2019). GSEAs were performed using GSEABase (version 1.48.0) and gene sets from the MSigDB collections (version 6.2) (Liberzon et al., 2015; Liberzon et al., 2011; Subramanian et al., 2005). Venn diagram analyses were performed using the tool provided on the Bioinformatics & Evolutionary Genomics website of Ghent University (Van de Peer).

### Preparation of graphs

Graphs were generated using Metascape (Zhou et al., 2019), RStudio (version 1.1.456) and the GraphPad Prism software (v.9.2.0).

## ACKNOWLEDGMENTS

We thank Thomas Krucker and Anke Harupa-Chung for providing assistance and mentorship during the execution of this study. We thank Sam Hofman, Ivonne Nieuwenhuis and Nicole van der Werff for expert technical assistance. This work was supported by grants from the Bill and Melinda Gates Foundation (T.T.D., C.H.M.K., G.R.; Grant no. OPP1141292, and T.T.D.; Grant no. INV010720) and Wellcome Trust (T.T.D, C.H.M.K.; Grant no. WT078285 and WT096157).

## CONTRIBUTIONS

**Gabriel Mitchell**: Conceptualization, Investigation, Supervision, Formal Analysis, Visualization, Writing—original draft, Writing—review and editing

**Guglielmo Roma**: Conceptualization, Resources, Data curation, Supervision, Funding acquisition, Investigation, Methodology, Project administration, Writing—review and editing

**Annemarie Voorberg-van der Wel**: Investigation, Methodology, Writing—review and editing

**Martin Beibel**: Data curation, Formal analysis, Investigation, Methodology, Software, Writing—review and editing

**Anne-Marie Zeeman**: Investigation, Methodology, Writing—review and editing

**Sven Schuierer**: Data curation, Software, Investigation, Methodology, Writing—review and editing

**Laura Torres**: Investigation Erika L Flannery: Investigation

**Clemens HM Kocken**: Conceptualization, Resources, Supervision, Funding acquisition, Investigation, Project administration, Writing—review and editing

**Sebastian A. Mikolajczak**: Supervision, Investigation

**Thierry T. Diagana**: Conceptualization, Resources, Supervision, Funding acquisition, Investigation, Project administration

